# Specification of frequency criteria for secondary findings genes to improve variant classification concordance

**DOI:** 10.1101/2025.09.19.676605

**Authors:** Jennifer J. Johnston, Kristy Lee, Deborah I. Ritter, Steven M. Harrison, Leslie G. Biesecker

## Abstract

**Purpose:** The return of secondary findings (SF) is well-established in clinical testing and is increasingly employed in clinical research testing. Variant classification can be challenging due to the lack of monogenic disease entity (MDE, defined as a gene-phenotype pair) specific variant classification recommendations. A key criterion for variant classification is disease allele frequency thresholds (DAFTs), which are not established for all MDEs recommended for the return of SF.

**Methods:** We calculated DAFTs for SF MDEs considering prevalence, gene contribution, and penetrance. ACMG criterion BS1 was set at the calculated DAFT value, with BA1 set at ten times the calculated DAFT. For genes associated with multiple SF MDEs without clear genotype-phenotype correlation, DAFT values were combined. GnomAD Grpmax filtering allele frequencies (FAF) for pathogenic/likely pathogenic classified variants were compared to calculated thresholds.

**Results:** We determined BS1 and BA1 values for 58 SF MDEs (47 genes). No pathogenic/likely pathogenic variant Grpmax FAF was greater than the relevant MDE-specific BA1.

**Conclusion:** Setting BA1 and BS1 thresholds should improve variant classification consistency and reduce misclassifications. For SF MDEs without current ClinGen Variant Curation Expert Panel (VCEP) specifications, these frequency specifications can be used until full criteria are available.

## Introduction

In 2013, the American College of Medical Genetics and Genomics (ACMG) recommended that secondary findings (SF) from clinical exome and genome sequencing be offered to individuals. Toward standardizing this process, the ACMG has put forward a list of monogenic disease entities (MDEs, defined as a gene-phenotype pair) for variant return, which is now updated annually. This work covers 94 MDEs (in 83 genes) suggested for SF return (v3.3) [1]. In 2015 the ACMG and the Association for Molecular Pathology (AMP) put forward criteria for classifying the pathogenicity of sequence variants [2]. Subsequent work showed discordance in variant classification amongst laboratories using the ACMG/AMP classification criteria [3]. To increase concordance, ClinGen is providing additional guidance for applying these guidelines through the Variant Classification working group (VC WG) and Variant Curation Expert Panels (VCEPs) for specific MDEs (https://clinicalgenome.org/) [4]. To help laboratories determine which variants should be reportable as SF, ClinGen, in partnership with the ACMG Secondary Findings Maintenance Working Group (SFWG), is providing SF specific guidance (https://search.clinicalgenome.org/kb/genes/acmgsf). As part of this effort we set out to calculate maximum credible population disease allele frequencies thresholds (DAFT), and specify values for BA1 (the rule code associated with variants too common, stand-alone benign) and BS1 (the rule code associated with variants likely too common, strong benign evidence) for SF MDEs that do not yet have specific classification criteria from a ClinGen VCEP. Harmonizing the classification of Benign and Likely Benign evidence should reduce the reporting burden on testing laboratories and increase consistency across labs.

## METHODS

### VCEP Specifications

For SF MDEs with available ClinGen VCEP specifications, BA1 and BS1 values and other relevant data are available on the ClinGen Specification Registry (https://cspec.genome.network/). BA1 and BS1 values available in the ClinGen Specification Registry (as of June 15, 2025) are included in Table 1. Note that because *HFE*-related hemochromatosis is recommended for return of a single variant, it was not considered in this work. For MDEs in the planned scope of a ClinGen VCEP but without specifications available in the ClinGen Specification Registry, we consulted with the VCEP to determine if preliminary values for BA1 and BS1 could be provided for this project (Table 1).

**Table 1.** BA1 and BS1 provided by ClinGen Variant Curation Expert Panels.

| Monogenic Disease Entity | VCEP BA1<sup>a</sup> | VCEP BS1<sup>a</sup> | MANE Transcript | Variant Curation Expert Panel |
| --- | --- | --- | --- | --- |
| *ABCD1*-related adrenal leukodystrophy | 0.00017<sup>b</sup> | 0.000017<sup>b</sup> | NM_000033.4 | Peroxisomal Disorders |
| *ACTC1*-related hypertrophic cardiomyopathy | 0.001 | 0.0001 | NM_005159.5 | Cardiomyopathy |
| *ACVRL1*-related hereditary hemorrhagic telangiectasia | 0.01 | 0.002 | NM_000020.3 | Hereditary Hemorrhagic Telangiectasia |
| *APC*-related familial adenomatous polyposis | 0.001 | 0.00001 | NM_000038.6 | InSiGHT Hereditary Colorectal Cancer/Polypsis |
| *APOB*-related familial hypercholesterolemia | 0.005<sup>b</sup> | 0.002<sup>b</sup> | NM_000384.3 | Familial Hypercholesterolemia |
| *BMPR1A*-related juvenile polyposis syndrome | 0.0001<sup>b</sup> | 0.000001<sup>b</sup> | NM_004329.3 | InSiGHT Hereditary Colorectal Cancer/Polypsis |
| *BRCA1*-related breast and ovarian cancer syndrome | 0.001 | 0.0001 | NM_007294.4 | ENIGMA BRCA1 and BRCA2 |
| *BRCA2*-related breast and ovarian cancer syndrome | 0.001 | 0.0001 | NM_000059.4 | ENIGMA BRCA1 and BRCA2 |
| *CYP27A1*-related cerebrotendinous xanthomatosis | 0.0076<sup>b</sup> | 0.00076<sup>b</sup> | NM_000784.4 | Pediatric Cataract |
| *ENG*-related hereditary hemorrhagic telangiectasia | 0.01 | 0.002 | NM_001114753.3 | Hereditary Hemorrhagic Telangiectasia |
| *FBN1* -related Marfan syndrome | 0.001 | 0.00005 | NM_000138 | FBN1 |
| *GAA* -related Pompe disease | 0.01 | 0.005 | NM_000152.4 | Lysosomal Storage Disorders |
| *HNF1A*-related monogenic diabetes | 0.0001 | 0.000033 | NM_000545.8 | Monogenic Diabetes |
| *KCNQ1*-related long-QT syndrome | 0.004 | 0.0004 | NM_000218.3 | Potassium Channel Arrhythmia |
| *LDLR*-related familial hypercholesterolemia | 0.005 | 0.002 | NM_000527.5 | Familial Hypercholesterolemia |
| *MLH1*-related Lynch syndrome | 0.001 | 0.0001 | NM_000249.4 | InSiGHT Hereditary Colorectal Cancer/Polypsis |
| *MSH2*-related Lynch syndrome | 0.001 | 0.0001 | NM_000251.3 | InSiGHT Hereditary Colorectal Cancer/Polypsis |
| *MSH6*-related Lynch syndrome | 0.0022 | 0.00022 | NM_000179.3 | InSiGHT Hereditary Colorectal Cancer/Polypsis |
| *MUTYH*-related familial adenomatous polyposis | 0.01054<sup>b</sup> | 0.00105<sup>b</sup> | NM_001048174.2 | InSiGHT Hereditary Colorectal Cancer/Polypsis |
| *MYBPC3*-related hereditary cardiomyopathy | 0.001 | 0.0002 | NM_000256.3 | Cardiomyopathy |
| *MYH7*-related DCM/HCM combined | 0.001 | 0.0001 | NM_000257.4 | Cardiomyopathy |
| *MYL2*-related hypertrophic cardiomyopathy | 0.001 | 0.0001 | NM_000432.4 | Cardiomyopathy |
| *MYL3* -related hypertrophic cardiomyopathy | 0.001 | 0.0001 | NM_000258.3 | Cardiomyopathy |
| *OTC*-related ornithine transcarbamylase deficiency | 0.01<sup>b</sup> | 0.002<sup>b</sup> | NM_000531.6 | Urea Cycle Disorders |
| *PALB2*-related breast cancer predisposition | 0.001 | 0.0001 | NM_024675.3 | Hereditary Breast, Ovarian and Pancreatic Cancer |
| *PCSK9*-related familial hypercholesterolemia | 0.0002<sup>b</sup> | 0.0001<sup>b</sup> | NM_174936.4 | Familial Hypercholesterolemia |
| *PMS2*-related Lynch syndrome | 0.0028 | 0.0001 | NM_000535.7 | InSiGHT Hereditary Colorectal Cancer/Polypsis |
| *PTEN* -related hamartoma tumor syndrome | 0.00056 | 0.000043 | NM_000314.8 | PTEN |
| *RPE65*-related retinopathy | 0.008 | 0.0008 | NM_000329.3 | Leber Congenital Amaurosis/early onset Retinal Dystrophy |
| *RYR1*-related malignant hyperthermia | 0.0038 | 0.0008 | NM_000540.3 | Malignant Hyperthermia Susceptibility |
| *SMAD4*-related juvenile polyposis syndrome / hereditary hemorrhagic telangiectasia | 0.0001<sup>b</sup> | 0.000001<sup>b</sup> | NM_005359.6 | InSiGHT Hereditary Colorectal Cancer/Polypsis |
| *TNNI3*-related hypertrophic cardiomyopathy | 0.001 | 0.0001 | NM_000363.5 | Cardiomyopathy |
| *TNNT2*-related DCM/HCM | 0.001 | 0.0001 | NM_001276345.2 | Cardiomyopathy |
| *TP53* -related Li-Fraumeni syndrome | 0.001 | 0.0003 | NM_000546.6 | TP53 |
| *TPM1*-related hypertrophic cardiomyopathy | 0.001 | 0.0001 | NM_001018005.2 | Cardiomyopathy |
| *VHL* -related von Hippel-Lindau syndrome | 0.000156 | 0.0000156 | NM_000551.4 | VHL |

DCM, Dilated cardiomyopathy; LQTS, Long QT syndrome; HCM, Hypertrophic cardiomyopathy.
<sup>a</sup>Some ClinGen VCEPs have recommendations for BS1\_supporting as well as additional instructions for BA1/BS1 use. Please access the ClinGen Criteria Specification Registry for the most complete and up to date information. <sup>b</sup>Preliminary ClinGen VCEP specifications not yet approved.

### BA1/BS1

For 58 MDEs included on the ACMG SF list without available BA1/BS1 values from a ClinGen VCEP we first established a DAFT for pathogenic variants. In theory, the DAFT could be calculated using the sum of all pathogenic variant allele frequencies across each MDE for the gene. The DAFT can be estimated using the MDE prevalence and penetrance. When a heritable phenotype has locus heterogeneity, however, MDE prevalence can be difficult to estimate and one approach is to estimate the combined MDE prevalence for the condition and the individual gene contribution separately. Values for condition prevalence, gene contribution, and penetrance were identified from the following resources: GeneReviews, the ClinGen Actionability Working Groups (https://clinicalgenome.org/working-groups/actionability/), and published literature (Table S1).

The MDE-specific DAFTs were calculated using the equations presented below. The allele frequency calculator on cardiodb.org [5] can be used to calculate the DAFT for autosomal monoallelic and biallelic MDEs with the “allelic heterogeneity” slider set to 1.

The following equation was used to calculate DAFT for MDEs expected to be inherited in an autosomal dominant pattern:

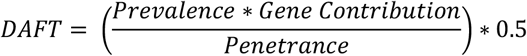

For MDEs expected to be inherited in an autosomal recessive pattern the following equation was used to determine DAFT:

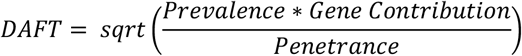

For MDEs inherited in an X-linked recessive pattern the following equation was used to determine DAFT:

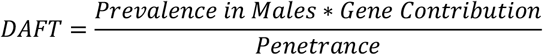

When considering SF variant classification for genes with multiple SF MDEs, genotype-phenotype correlation was taken into consideration to determine a combined DAFT for the gene. For genes where variants demonstrate strong correlation to an allelic MDE (i.e., loss of function variants in *SCN5A* (HGNC:10593) associated with Brugada syndrome and gain of function variants in *SCN5A* associated with Long QT syndrome (LQTS)), DAFT values were calculated for each SF MDE individually and the highest DAFT value was used. For genes where variants do not demonstrate strong correlation to a specific SF MDE (i.e., variants in *DSP* (HGNC: 3052) are associated with either arrhythmogenic right ventricular cardiomyopathy or dilated cardiomyopathy), DAFT values were calculated for each SF MDE individually and then summed. For genes where variants are associated with non-SF MDEs and SF MDEs (i.e., variants in *RYR1* (HGNC: 10483) are associated with malignant hyperthermia and myopathy), only DAFTs for the SF MDEs (malignant hyperthermia) are considered. When possible ClinGen VCEPs were consulted for input on setting these values.

The calculated DAFT values were used to generate BA1 and BS1 values with the goal of setting thresholds that would err on the side of being (numerically) too high, as this would minimize the chance that the variants are incorrectly classified as benign. The BA1 and BS1 values were set at ten times and one times the DAFT, respectively. This was done to minimize the error of a numerically low threshold inappropriately filtering out variants from full classification (BA1) or erroneously applying benign evidence (BS1).

### Validation of BA1/BS1

Calculated values for BA1 and BS1 were compared to gnomAD v4.1 continental group maximum (Grpmax) filtering allele frequencies (FAF) for the most common variants classified as pathogenic or likely pathogenic in ClinVar (P/LP variants, classifications from ClinVar as provided in gnomAD v4.1). ClinVar classification of pathogenicity was reviewed for P/LP variants with a Grpmax FAF > DAFT. For genes where loss of function is the mechanism for the SF MDE, Grpmax FAFs for putative loss of function (pLoF) variants in the MANE Select transcript, but not present in ClinVar, were also considered (novel pLoF flagged as low confidence pLoF in gnomAD were excluded). For *TTN* (HGNC:12403), pLoF variants were only considered in the A band [6].

## RESULTS

Twenty-eight MDEs (not considering *HFE*-related hemochromatosis) have ClinGen VCEP specifications available and preliminary specifications were provided by VCEPs for an additional eight MDEs (Table1, Supplemental Table 2). Fifty-eight MDEs (47 genes), lack available VCEP specifications, seventeen of which are awaiting specifications by a VCEP and 41 do not have a currently active ClinGen VCEP. For these 58 SF MDEs, we calculated DAFT values using the provided equations and used these to determine values for BA1 and BS1 (Tables 2 and 3, Supplemental Table 2). These DAFT/BA1/BS1 values are only relevant for the SF MDE as listed and should not be used for other MDEs.

**Table 2.**
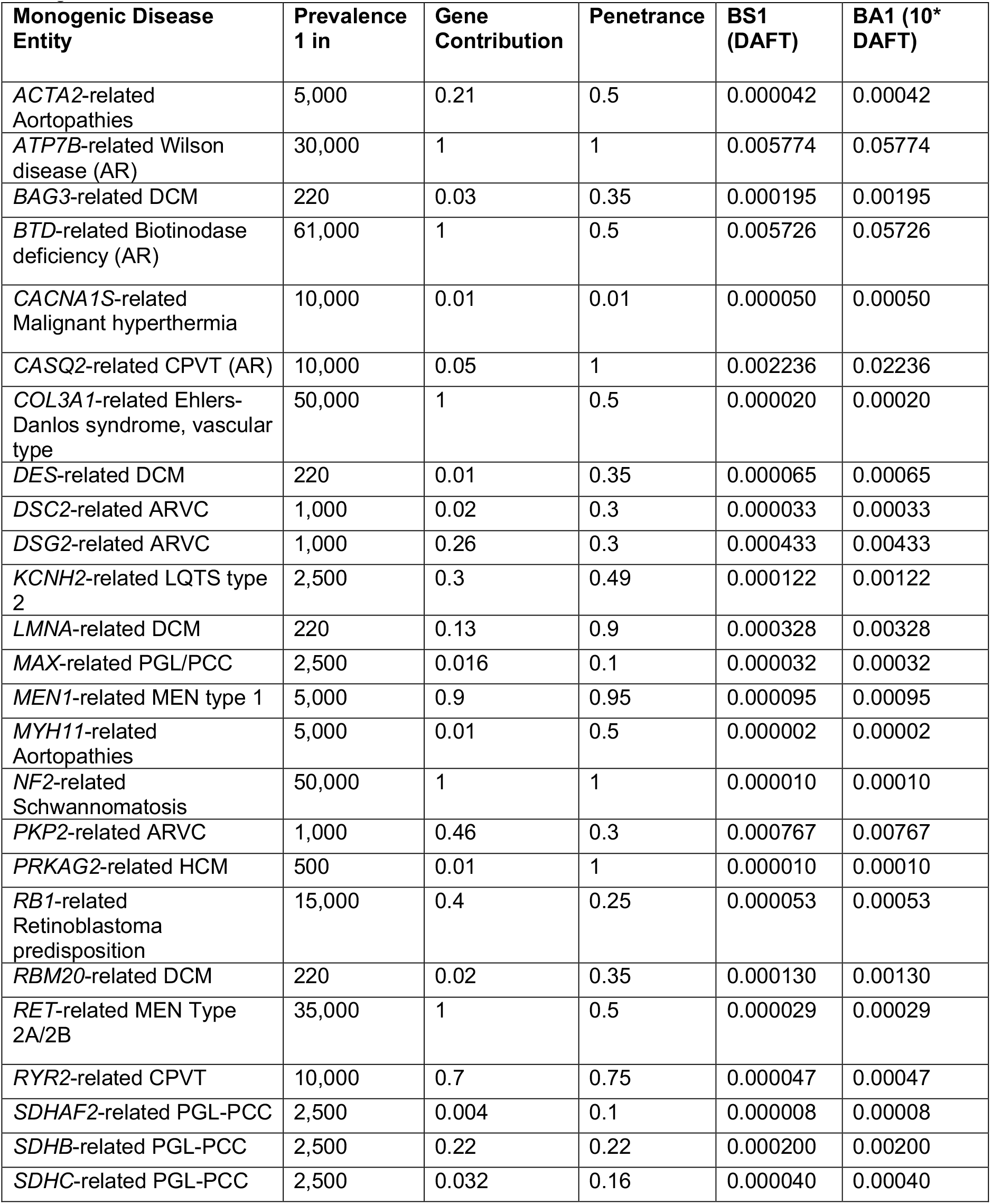
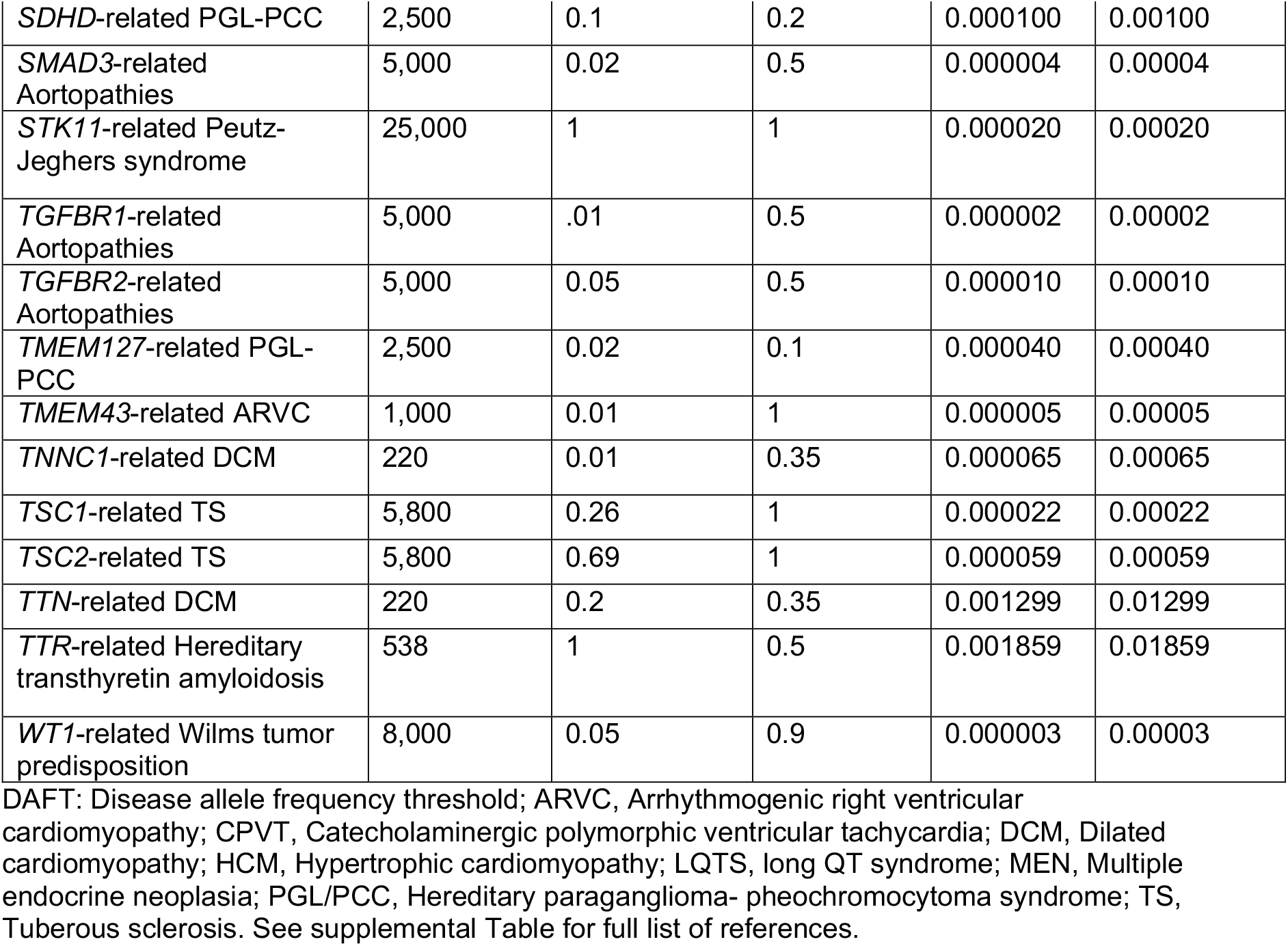
BA1 and BS1 values calculated for genes reported for a single secondary finding monogenic disease entities.

| **Monogenic Disease Entity** | **Prevalence 1 in** | **Gene Contribution** | **Penetrance** | **BS1 (DAFT)** | **BA1 (10\* DAFT)** |
| --- | --- | --- | --- | --- | --- |
| *ACTA2*-related Aortopathies | 5,000 | 0.21 | 0.5 | 0.000042 | 0.00042 |
| *ATP7B*-related Wilson disease (AR) | 30,000 | 1 | 1 | 0.005774 | 0.05774 |
| *BAG3*-related DCM | 220 | 0.03 | 0.35 | 0.000195 | 0.00195 |
| *BTD*-related Biotinodase deficiency (AR) | 61,000 | 1 | 0.5 | 0.005726 | 0.05726 |
| *CACNA1S*-related Malignant hyperthermia | 10,000 | 0.01 | 0.01 | 0.000050 | 0.00050 |
| *CASQ2*-related CPVT (AR) | 10,000 | 0.05 | 1 | 0.002236 | 0.02236 |
| *COL3A1*-related Ehlers-Danlos syndrome, vascular type | 50,000 | 1 | 0.5 | 0.000020 | 0.00020 |
| *DES*-related DCM | 220 | 0.01 | 0.35 | 0.000065 | 0.00065 |
| *DSC2*-related ARVC | 1,000 | 0.02 | 0.3 | 0.000033 | 0.00033 |
| *DSG2*-related ARVC | 1,000 | 0.26 | 0.3 | 0.000433 | 0.00433 |
| *KCNH2*-related LQTS type 2 | 2,500 | 0.3 | 0.49 | 0.000122 | 0.00122 |
| *LMNA*-related DCM | 220 | 0.13 | 0.9 | 0.000328 | 0.00328 |
| *MAX*-related PGL/PCC | 2,500 | 0.016 | 0.1 | 0.000032 | 0.00032 |
| *MEN1*-related MEN type 1 | 5,000 | 0.9 | 0.95 | 0.000095 | 0.00095 |
| *MYH11*-related Aortopathies | 5,000 | 0.01 | 0.5 | 0.000002 | 0.00002 |
| *NF2*-related Schwannomatosis | 50,000 | 1 | 1 | 0.000010 | 0.00010 |
| *PKP2*-related ARVC | 1,000 | 0.46 | 0.3 | 0.000767 | 0.00767 |
| *PRKAG2*-related HCM | 500 | 0.01 | 1 | 0.000010 | 0.00010 |
| *RB1*-related Retinoblastoma predisposition | 15,000 | 0.4 | 0.25 | 0.000053 | 0.00053 |
| *RBM20*-related DCM | 220 | 0.02 | 0.35 | 0.000130 | 0.00130 |
| *RET*-related MEN Type 2A/2B | 35,000 | 1 | 0.5 | 0.000029 | 0.00029 |
| *RYR2*-related CPVT | 10,000 | 0.7 | 0.75 | 0.000047 | 0.00047 |
| *SDHAF2*-related PGL-PCC | 2,500 | 0.004 | 0.1 | 0.000008 | 0.00008 |
| *SDHB*-related PGL-PCC | 2,500 | 0.22 | 0.22 | 0.000200 | 0.00200 |
| *SDHC*-related PGL-PCC | 2,500 | 0.032 | 0.16 | 0.000040 | 0.00040 |
| *SDHD*-related PGL-PCC | 2,500 | 0.1 | 0.2 | 0.000100 | 0.00100 |
| *SMAD3*-related Aortopathies | 5,000 | 0.02 | 0.5 | 0.000004 | 0.00004 |
| *STK11*-related Peutz-Jeghers syndrome | 25,000 | 1 | 1 | 0.000020 | 0.00020 |
| *TGFBR1*-related Aortopathies | 5,000 | .01 | 0.5 | 0.000002 | 0.00002 |
| *TGFBR2*-related Aortopathies | 5,000 | 0.05 | 0.5 | 0.000010 | 0.00010 |
| *TMEM127*-related PGL-PCC | 2,500 | 0.02 | 0.1 | 0.000040 | 0.00040 |
| *TMEM43*-related ARVC | 1,000 | 0.01 | 1 | 0.000005 | 0.00005 |
| *TNNC1*-related DCM | 220 | 0.01 | 0.35 | 0.000065 | 0.00065 |
| *TSC1*-related TS | 5,800 | 0.26 | 1 | 0.000022 | 0.00022 |
| *TSC2*-related TS | 5,800 | 0.69 | 1 | 0.000059 | 0.00059 |
| *TTN*-related DCM | 220 | 0.2 | 0.35 | 0.001299 | 0.01299 |
| *TTR*-related Hereditary transthyretin amyloidosis | 538 | 1 | 0.5 | 0.001859 | 0.01859 |
| *WT1*-related Wilms tumor predisposition | 8,000 | 0.05 | 0.9 | 0.000003 | 0.00003 |

DAFT: Disease allele frequency threshold; ARVC, Arrhythmogenic right ventricular cardiomyopathy; CPVT, Catecholaminergic polymorphic ventricular tachycardia; DCM, Dilated cardiomyopathy; HCM, Hypertrophic cardiomyopathy; LQTS, long QT syndrome; MEN, Multiple endocrine neoplasia; PGL/PCC, Hereditary paraganglioma- pheochromocytoma syndrome; TS, Tuberous sclerosis. See supplemental Table for full list of references.

**Table 3.** BA1 and BS1 values calculated for genes reported for multiple secondary finding monogenic disease entities.

| **Monogenic Disease** | **Prevalence**<br>**1 in** | **Gene**<br>**Contribution** | **Penetrance** | **BS1**<br>**(DAFT)** | **BA1**<br>**(10\* DAFT)** |
| --- | --- | --- | --- | --- | --- |
| *CALM1*-related LQTS | 2,500 | 0.01 | 0.4 | 0.000005 | 0.00005 |
| *CALM1*-related CPVT | 10,000 | 0.01 | 0.63 | 0.000001 | 0.00001 |
| *CALM1* COMBINED<br>VALUES |  |  |  | 0.000006 | 0.00006 |
| *CALM2*-related LQTS | 2,500 | 0.01 | 0.4 | 0.000005 | 0.00005 |
| *CALM2*-related CPVT | 10,000 | 0.01 | 0.63 | 0.000001 | 0.00001 |
| *CALM2* COMBINED<br>VALUES |  |  |  | 0.000006 | 0.00006 |
| *CALM3*-related LQTS | 2,500 | 0.01 | 0.4 | 0.000005 | 0.00005 |
| *CALM3*-related CPVT | 10,000 | 0.01 | 0.63 | 0.000001 | 0.00001 |
| *CALM3* COMBINED<br>VALUES |  |  |  | 0.000006 | 0.00006 |
| *DSP*-related ARVC | 1,000 | 0.39 | 0.3 | 0.000650 | 0.00650 |
| *DSP*-related DCM | 220 | 0.02 | 0.35 | 0.000130 | 0.00130 |
| *DSP* COMBINED<br>VALUES |  |  |  | 0.000780 | 0.00780 |
| *FLNC*-related DCM | 220 | 0.04 | 0.35 | 0.000260 | 0.00260 |
| *FLNC*-related HCM | 500 | 0.02 | 0.75 | 0.000027 | 0.00027 |
| *FLNC* COMBINED<br>VALUES |  |  |  | 0.000286 | 0.00286 |
| *GLA*-related Fabry<br>disease | 3,100 | 1 | 1 | 0.000323 | 0.00323 |
| *GLA*-related HCM | 500 | 0.01 | .5 | 0.000040 | 0.00040 |
| *GLA* COMBINED<br>VALUES |  |  |  | 0.000363 | 0.003626 |
| PLN-related ARVC | 1,000 | 0.01 | 0.3 | 0.000017 | 0.00017 |
| PLN-related HCM | 500 | 0.03 | 0.5 | 0.000060 | 0.00060 |
| PLN-related DCM | 220 | 0.01 | 0.35 | 0.000065 | 0.00065 |
| PLN COMBINED<br>VALUES | NA | NA | NA | 0.000142 | 0.00142 |
| *SCN5A*-related Brugada<br>syndrome | 830 | 0.3 | 0.2 | 0.000904 | 0.00904 |
| *SCN5A*-related LQTS | 2,500 | 0.10 | 0.42 | 0.000190 | 0.00190 |
| *SCN5A*-related DCM | 220 | 0.02 | 0.35 | 0.000130 | 0.00130 |
| *SCN5A* COMBINED<br>VALUES<br>Brugada and DCM |  |  |  | 0.00103 | 0.01033 |
| *TRDN*-related CPVT (AR) | 10,000 | 0.02 | 0.75 | 0.001633 | 0.016329932 |
| *TRDN*-related LQTS (AR) | 2,500 | 0.01 | 0.41 | 0.003123 | 0.03123 |
| *TRDN* COMBINED<br>VALUES (AR) |  |  |  | 0.004756 | 0.04756 |

DAFT: Disease allele frequency threshold; AR, Autosomal recessive; ARVC, Arrhythmogenic right ventricular cardiomyopathy; CPVT, Catecholaminergic polymorphic ventricular tachycardia; DCM, Dilated cardiomyopathy; LQTS, Long QT syndrome; HCM, Hypertrophic cardiomyopathy. See supplemental table for full list of references.

For nine genes included in this study, multiple MDEs are recommended for SF return. For eight of these genes, a clear genotype/phenotype correlation was not identified, and DAFT values were combined across the reportable allelic MDEs (Table 3). Variants in *SCN5A* are recommended for return for three distinct MDEs. Loss of function variants are associated with Brugada syndrome and gain of function variants are associated with LQTS. *SCN5A* variants are also associated with DCM, but without a clear genotype/phenotype correlation to distinguish them from the Brugada-or LQTS-associated variants. For *SCN5A*, the DAFTs for Brugada and LQTS were determined and the greater of the two was combined with the DAFT for DCM.

As the goal of this work was to set numerically conservative values for BA1/BS1 we compared gnomAD Grpmax FAF for ClinVar P/LP and novel pLoF variants to the calculated values for BA1/BS1. With regards to BA1 thresholds, we identified no P/LP or pLoF variant (in genes where loss of function is a mechanism for the SF MDEs) with a gnomAD Grpmax FAF greater than the calculated BA1 value for the relevant gene. With regards to BS1 thresholds, for 39 genes no P/LP or pLoF variant was identified with a Grpmax FAF higher than the calculated BS1 value. A total of nine P/LP or pLoF variants in eight genes had a Grpmax FAF greater than the calculated BS1 value for the relevant gene (Table 4). These nine variants represent 0.2% of a total of 4,305 P/LP or pLoF variants.

**Table 4.**
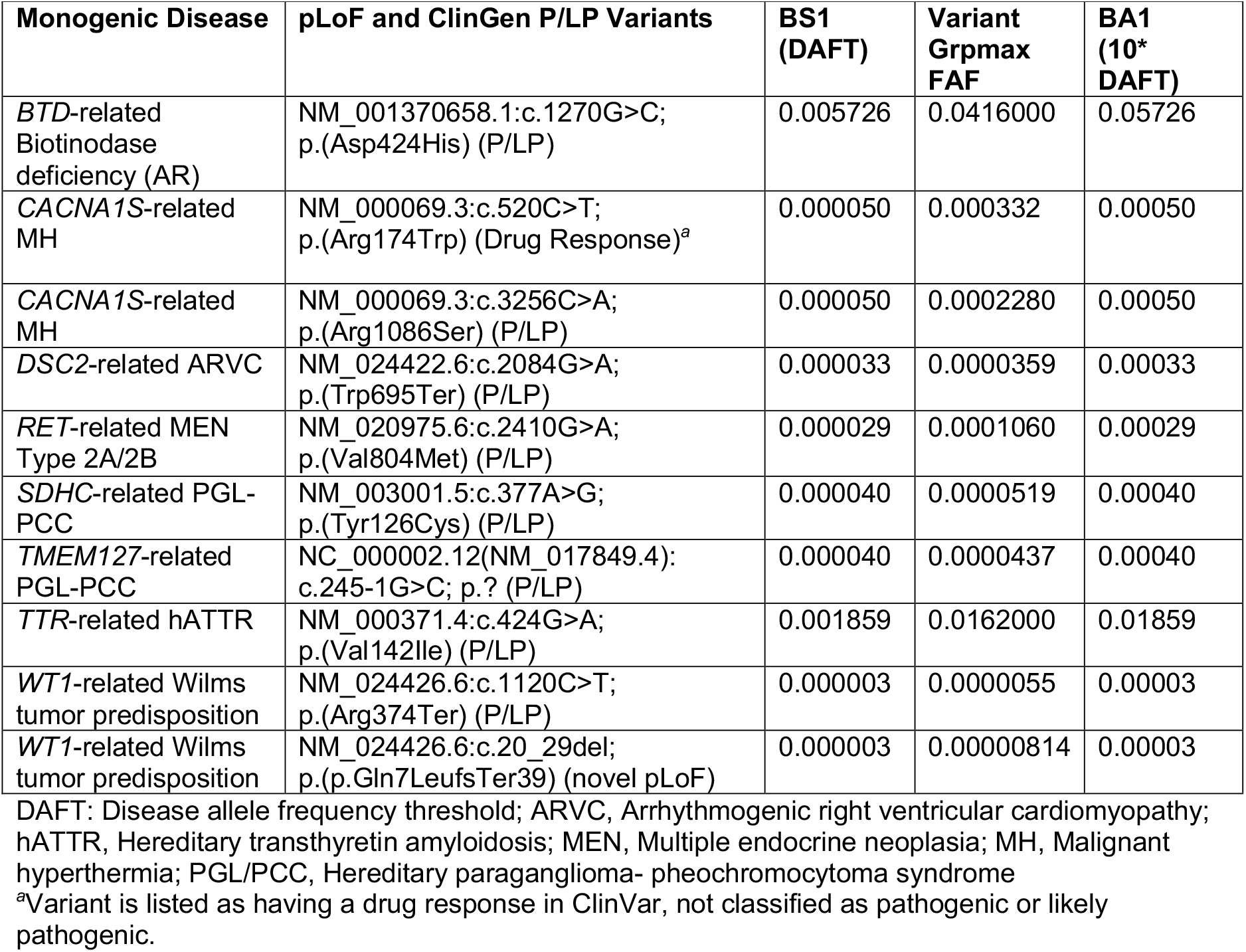
Variants with gnomAD Grpmax filtering allele frequencies greater than calculated DAFT.

| Monogenic Disease | pLoF and ClinGen P/LP Variants | BS1 (DAFT) | Variant Grpmax FAF | BA1 (10* DAFT) |
| --- | --- | --- | --- | --- |
| *BTD*-related Biotinodase deficiency (AR) | NM_001370658.1:c.1270G>C; p.(Asp424His) (P/LP) | 0.005726 | 0.0416000 | 0.05726 |
| *CACNA1S*-related MH | NM_000069.3:c.520C>T; p.(Arg174Trp) (Drug Response)<sup>a</sup> | 0.000050 | 0.000332 | 0.00050 |
| *CACNA1S*-related MH | NM_000069.3:c.3256C>A; p.(Arg1086Ser) (P/LP) | 0.000050 | 0.0002280 | 0.00050 |
| *DSC2*-related ARVC | NM_024422.6:c.2084G>A; p.(Trp695Ter) (P/LP) | 0.000033 | 0.0000359 | 0.00033 |
| *RET*-related MEN Type 2A/2B | NM_020975.6:c.2410G>A; p.(Val804Met) (P/LP) | 0.000029 | 0.0001060 | 0.00029 |
| *SDHC*-related PGL-PCC | NM_003001.5:c.377A>G; p.(Tyr126Cys) (P/LP) | 0.000040 | 0.0000519 | 0.00040 |
| *TMEM127*-related PGL-PCC | NC_000002.12(NM_017849.4): c.245-1G>C; p.? (P/LP) | 0.000040 | 0.0000437 | 0.00040 |
| *TTR*-related hATTR | NM_000371.4:c.424G>A; p.(Val142Ile) (P/LP) | 0.001859 | 0.0162000 | 0.01859 |
| *WT1*-related Wilms tumor predisposition | NM_024426.6:c.1120C>T; p.(Arg374Ter) (P/LP) | 0.000003 | 0.0000055 | 0.00003 |
| *WT1*-related Wilms tumor predisposition | NM_024426.6:c.20_29del; p.(p.Gln7LeufsTer39) (novel pLoF) | 0.000003 | 0.00000814 | 0.00003 |

DAFT: Disease allele frequency threshold; ARVC, Arrhythmogenic right ventricular cardiomyopathy; hATTR, Hereditary transthyretin amyloidosis; MEN, Multiple endocrine neoplasia; MH, Malignant hyperthermia; PGL/PCC, Hereditary paraganglioma- pheochromocytoma syndrome
<sup>a</sup>Variant is listed as having a drug response in ClinVar, not classified as pathogenic or likely pathogenic.

## Discussion

Variant classification using the ACMG criteria is facilitated when gene-specific criteria for MDEs are applied. In the 2015 ACMG/AMP classification criteria, a Grpmax minor allele frequency (MAF) threshold of 0.05 was set as a stand-alone benign criterion (BA1). Application of BA1 allowed a variant with a Grpmax MAF greater than 0.05 to be classified as benign without further analysis. It was later recognized that a few common pathogenic variants had a Grpmax MAF greater than 0.05, requiring caution to be taken when using this criterion [7]. It is important to note that if the BA1 threshold is set inappropriately low for an MDE, a variant that deserves full classification might be filtered out and not further considered. However, for many MDEs the most common pathogenic variant has a Grpmax MAF in gnomAD orders of magnitude below 0.05 and setting BA1 at 0.05 substantially increases the analysis burden of variants that are too common to be pathogenic for that MDE. An MDE-specific BA1 can decrease the variant set requiring full classification and protect against pathogenic variants being incorrectly classified as benign based on frequency alone. For the set of SF MDEs considered in this study, setting BA1 at ten times the DAFT allowed for filtering of common variants while retaining P/LP variants for full classification. Of note, we did not consider allelic heterogeneity when setting MDE-specific BA1 values. For MDE with many rare pathogenic variants the resulting BA1 may still be orders of magnitude higher than the Grpmax FAF for P/LP variants. When VCEPs determine gene-specific criteria for MDEs, allelic heterogeneity should be considered.

In addition to BA1, we set values for BS1, to allow for population frequency to be used as strong evidence against pathogenicity. An appropriate value for BS1 would minimize false negatives (true pathogenic variants awarded BS1) while providing strong benign evidence to variants that are more common than expected for an MDE. Unlike BA1, BS1 does not negate full variant classification, and the expectation is that BS1 will be set such that there is a small probability that a pathogenic variant will be awarded BS1. For the set of SF MDE included in this study nine out of 4,305 P/LP or pLoF variants in gnomAD, or 0.2%, had a Grpmax FAF that allowed for the use of BS1 (Table 4). In theory, if a pathogenic variant has a Grpmax FAF greater than BS1, which is equal to the DAFT, then the DAFT must be incorrect since the DAFT should account for all pathogenic variants. However, to calculate the DAFT one must use values for MDE prevalence, gene contribution, and penetrance that are often not well defined. Conservative values (highest reasonable MDE prevalence and gene contribution and lowest reasonable penetrance) were used in this work to err on the side of setting a higher, and thus “safer” DAFT (less variants meet BA1/BS1 and more variants need full classification).

Inheritance patterns as defined for ACMG SF reporting were applied in this work and should be considered when using the calculated values for variant classification. *DSC2*-related arrhythmogenic right ventricular cardiomyopathy (ARVC) is thought to generally be inherited in an autosomal dominant pattern and monoallelic inheritance is considered for ACMG SF reporting of *DSC2*-related ARVC. Some variants in *DSC2* (HGNC:3036), however, including the loss of function variant identified in the Hutterite population, (NM_024422.6:c.1660C>T, p.(Gln554Ter)), have been reported to only cause ARVC when biallelic [8, 9]. The *DSC2* P/LP variant identified here (NM_024422.6:c.2084G>A, p.(Trp695Ter)) with a Grpmax FAF (0.000036) greater than the BS1 threshold (0.000033) has not been reported to our knowledge in heterozygosity in an affected individual and may instead be associated with biallelic (autosomal recessive) inheritance of ARVC. Variants associated with biallelic inheritance for ARVC are not considered reportable as a SF when identified in carriers. A biallelic inheritance model would result in a higher DAFT as compared to using a monoallelic inheritance model making it imperative that the inheritance model is considered when applying BA1/BS1.

For some disorders, including *CACNA1S*-related malignant hyperthermia (MH), setting an appropriate DAFT is complicated by disease attributes. An MH diagnosis is made when an individual has a reaction to inhaled anesthetics and since not all individuals are exposed to inhaled anesthetics, and one exposure may not trigger symptoms, it is very difficult to know the true prevalence of the disorder and the true penetrance. Here we have used the values for disease prevalence and penetrance as determined by the MH VCEP for *RYR1* as best estimates [10]. An additional complicating factor for setting an accurate DAFT for *CACNA1S*-related MH, is that only a few P/LP variants have been identified so the true DAFT may be very close to the actual variant frequencies. One might consider increasing the BS1 value when few P/LP variants have been identified in a gene.

Even when using well-defined values for MDE prevalence, penetrance, and gene contribution to calculate the DAFT, variant-specific reduced penetrance can complicate the generalizability of our proposed BA1/BS1 thresholds. Variant-specific reduced penetrance may cause a variant to have a higher Grpmax FAF than would be expected based on the attributes of the relevant MDE. The NM_020975.6:c.2410G>A; p.(Val804Met) *RET* (HGNC:9967) variant has reduced penetrance compared to other variants that contribute to *RET*-related medullary thyroid cancer [11] and the Grpmax FAF is approximately four times the relevant BS1 threshold. The NM_000371.4:c.424G>A; p.(Val142Ile) variant in *TTR* (HGNC:12405) is thought to have reduced penetrance compared to other variants that contribute to *TTR*-related hereditary transthyretin amyloidosis and the Grpmax FAF is approximately nine times the relevant BS1. This variant is common in individuals of West African descent and while individuals with this variant have been shown to have transthyretin amyloid deposits by 65 years of age, the occurrence of a recognizable phenotype in these individuals is reduced [12]. While reports of reduced penetrance for the NM_003001.5:c.377A>G; p.(Tyr126Cys) variant in *SDHC* (HGNC:10682) were not identified, this variant is relatively rare in pheochromocytoma/paraganglioma cohorts but has the highest gnomAD Grpmax FAF for ClinVar *SDHC* P/LP variants [13]. Variant frequency in affected cohorts is expected to reflect the frequency in the corresponding continental population, however, this relationship can be disrupted by variant-specific penetrance. Hypomorphic alleles can also have a Grpmax FAF higher than expected for the relevant MDE. The hypomorphic NM_001370658.1:c.1270C>G; p.(Asp424His) variant in *BTD* (HGNC:1122) causes partial biotinidase deficiency when in *trans* to a second pathogenic variant [14]. Individuals who are homozygous for p.(Asp424His), however, do not demonstrate biotinidase deficiency allowing for an FAF higher than the calculated DAFT [15]. When applying BS1 the variant scientist needs to consider potential variant specific complicating factors including reduced penetrance and hypomorphic function.

The BS1 values in this study ranged from 0.000003 for *WT1*-related Wilms tumor to 0.0058 for *ATP7B*-related Wilson disease. The smaller the MDE-specific BS1 value the fewer alleles required in gnomAD to trigger its use and the more likely that random sampling error can lead to a FAF greater than BS1 by chance. For *WT1* (HGNC:12796), two P/LP variants met the relevant BS1 with fewer than five alleles identified in gnomAD. Consideration might be given to requiring a minimum number of alleles for triggering BS1.

ClinGen VCEPs combine the expertise of geneticists, physician specialists, laboratory geneticists, genetic counselors and basic scientists to consider the genetics, biology, and the molecular diagnostic approach of the MDE to derive variant classification recommendations, specifically, adaptations of ACMG/AMP 2015 general recommendations. Few laboratories have the expertise or resources to set MDE-specific criteria for all SF genes. In the absence of VCEP specifications, one can rely on the original ACMG criteria with ClinGen Variant Classification working group (VC WG) recommendations. However, for criteria such as BA1, the VC WG does not set thresholds independently of VCEPs, leaving many laboratories to default to the general 5% recommendation from the ACMG/AMP recommendations [2], which increases the analysis burden, or developing in house estimated MAF that increase discordance between laboratories’ variant classifications. We suggest a middle ground of setting numerically high values for BA1 and BS1, based on empirical data from DAFT calculations, sufficient to decrease the classification workload. In doing so, we deliberately err on the side of reduced filtering of variants that are benign (BA1) and missing some opportunities to add to evidence of benignity (BS1). When MDE-specific recommendations are not available, the laboratory geneticist is left to adapt the criteria for each variant, potentially leading to lower consistency across laboratories. We provide calculated SF MDE BA1/BS1 values with supporting prevalence, penetrance and gene contribution information as a resource for the community and plan to make them available on the ClinGen website (https://search.clinicalgenome.org/kb/genes/acmgsf). Importantly, VCEP specifications, when available will supersede this work. While full VCEP specifications are ideal, a set of available and consistent thresholds extracted from available published literature could improve accuracy and consistency across SF MDEs until they are developed for all genes.

## Acknowledgements

We thank the following ClinGen VCEPs for providing preliminary BA1/BS1 values: Familial Hypercholesterolemia VCEP, InSiGHT Hereditary Colorectal Cancer/Polyposis VCEP, Pediatric Cataract VCEP, Peroxisomal Disorders VCEP, Urea Cycle Disorders VCEP. The following VCEPs provided helpful discussions around appropriate MDE prevalence, penetrance, and gene contributions: Desmosomal Cardiomyopathy VCEP, Endocrine Tumor Predisposition VCEP, InSight Hereditary Colorectal Cancer/Polyposis VCEP, Potassium Channel Arrhythmia VCEP. We thank the ClinGen Variant Classification Working Group for helpful comments and discussion.

## Author Contributions

Conceptualization: JJJ, LGB; Data Curation: JJJ; Funding acquisition: LGB; Investigation: JJJ; Methodology: JJJ; Project Administration: JJJ; Resources: LGB; Supervision: JJJ; Writing-original draft: JJJ; Writing-review & editing: JJJ, KL, DIR, SMH, LGB

## Ethics Declaration

Not applicable

## Data Availability

Data is available upon individual request.

## REFERENCES

1. Lee, K, Abul-Husn, NS, Amendola, LM, et al., ACMG SF v3.3 list for reporting of secondary findings in clinical exome and genome sequencing: A policy statement of the American College of Medical Genetics and Genomics (ACMG). Genet Med, 2025. 27(8): p. 101454. 10.1016/j.gim.2025.101454

2. Richards, S, Aziz, N, Bale, S, et al., Standards and guidelines for the interpretation of sequence variants: a joint consensus recommendation of the American College of Medical Genetics and Genomics and the Association for Molecular Pathology. Genet Med, 2015. 17(5): p. 405–24. 10.1038/gim.2015.30

3. Amendola, LM, Muenzen, K, Biesecker, LG, et al., Variant Classification Concordance using the ACMG-AMP Variant Interpretation Guidelines across Nine Genomic Implementation Research Studies. Am J Hum Genet, 2020. 107(5): p. 932–941. 10.1016/j.ajhg.2020.09.011

4. Rehm, HL, Berg, JS, Brooks, LD, et al., ClinGen--the Clinical Genome Resource. N Engl J Med, 2015. 372(23): p. 2235–42. 10.1056/NEJMsr1406261

5. Whiffin, N, Minikel, E, Walsh, R, et al., Using high-resolution variant frequencies to empower clinical genome interpretation. Genet Med, 2017. 19(10): p. 1151–1158. 10.1038/gim.2017.26

6. Morales, A, Kinnamon, DD, Jordan, E, et al., Variant Interpretation for Dilated Cardiomyopathy: Refinement of the American College of Medical Genetics and Genomics/ClinGen Guidelines for the DCM Precision Medicine Study. Circ Genom Precis Med, 2020. 13(2): p. e002480. 10.1161/CIRCGEN.119.002480

7. Ghosh, R, Harrison, SM, Rehm, HL, et al., Updated recommendation for the benign stand-alone ACMG/AMP criterion. Hum Mutat, 2018. 39(11): p. 1525–1530. 10.1002/humu.23642

8. Chen, L, Hu, Y, Saguner, AM, et al., Natural History and Clinical Outcomes of Patients With DSG2/DSC2 Variant-Related Arrhythmogenic Right Ventricular Cardiomyopathy. Circulation, 2025. 151(17): p. 1213–1230. 10.1161/CIRCULATIONAHA.124.072226

9. Wong, JA, Duff, HJ, Yuen, T, et al., Phenotypic analysis of arrhythmogenic cardiomyopathy in the Hutterite population: role of electrocardiogram in identifying high-risk desmocollin-2 carriers. J Am Heart Assoc, 2014. 3(6): p. e001407. 10.1161/JAHA.114.001407

10. Johnston, JJ, Dirksen, RT, Girard, T, et al., Variant curation expert panel recommendations for RYR1 pathogenicity classifications in malignant hyperthermia susceptibility. Genet Med, 2021. 23(7): p. 1288–1295. 10.1038/s41436-021-01125-w

11. Loveday, C, Josephs, K, Chubb, D, et al., p.Val804Met, the Most Frequent Pathogenic Mutation in RET, Confers a Very Low Lifetime Risk of Medullary Thyroid Cancer. J Clin Endocrinol Metab, 2018. 103(11): p. 4275–4282. 10.1210/jc.2017-02529

12. Buxbaum, JN and Ruberg, FL, Transthyretin V122I (pV142I)* cardiac amyloidosis: an age-dependent autosomal dominant cardiomyopathy too common to be overlooked as a cause of significant heart disease in elderly African Americans. Genet Med, 2017. 19(7): p. 733–742. 10.1038/gim.2016.200

13. Williams, ST, Chatzikyriakou, P, Carroll, PV, et al., SDHC phaeochromocytoma and paraganglioma: A UK-wide case series. Clin Endocrinol (Oxf), 2022. 96(4): p. 499–512. 10.1111/cen.14594

14. Swango, KL, Demirkol, M, Huner, G, et al., Partial biotinidase deficiency is usually due to the D444H mutation in the biotinidase gene. Hum Genet, 1998. 102(5): p. 571–5. 10.1007/s004390050742

15. Pindolia, K, Jordan, M, and Wolf, B, Analysis of mutations causing biotinidase deficiency. Hum Mutat, 2010. 31(9): p. 983–91. 10.1002/humu.21303

